# Multi-level cellular and functional annotation of single-cell transcriptomes

**DOI:** 10.1101/2022.03.13.484162

**Authors:** Nicholas Mikolajewicz, Kevin R. Brown, Jason Moffat, Hong Han

## Abstract

Single-cell RNA-sequencing (scRNA-seq) offers unprecedented insight into heterogenous biology, allowing for the interrogation of cellular populations and gene expression programs at single-cell resolution. Here, we introduce scPipeline, a single-cell analytic toolbox that offers modular workflows for multi-level cellular annotation and user-friendly analysis reports. Novel methods that are introduced to facilitate scRNA-seq annotation include: *(i*) co-dependency index (CDI)-based differential expression; *(ii*) cluster resolution optimization using a marker-specificity criterion; *(iii*) marker-based cell-type annotation with Miko scoring; and *(iv*) gene program discovery using scale-free shared nearest neighbor network (SSN) analysis. Our unsupervised and supervised procedures were validated using a diverse collection of scRNA-seq datasets and we provide illustrative examples of cellular and transcriptomic annotation of developmental and immunological scRNA-seq atlases. Overall, scPipeline provides a flexible computational framework for in-depth scRNA-seq analysis.

## Introduction

Single-cell RNA-sequencing (scRNA-seq) has facilitated the characterization of diverse cellular populations at an unprecedented resolution, with the evolution of high-throughput protocols now allowing the profiling of millions of cells in a single experiment. While experimental protocols such as SMART-seq2^1^, Drop-seq^2^, sci-RNA-seq3^3^ and commercial 10X genomics vary in approach and scale, gene expression matrices (gene-by-cell count) are ultimately generated and represent a common starting point for most downstream analyses.

The development of computational toolboxes like Seurat^4-7^, Scanpy^8^, and Cell Ranger (10X Genomics, commercial) facilitate scRNA-seq analyses broadly across a diverse array of research topics. These tools offer application-tailored functionalities, including data pre-processing, normalization, quality control (QC) and clustering analysis. However, comprehensive analyses still require a degree of computational expertise. With the more recent emergence of interactive and notebook-based analysis platforms, scRNA-seq analysis has become more accessible to users lacking high-level computational skills^9-11^. Despite the user-friendly interface offered by these platforms, difficulties can arise with custom-tailored analyses, or when data integration between different scRNA-seq platforms is required. To address these limitations, we have developed scPipeline, a report-based single-cell analytic toolbox. scPipeline is offered as a series of Rmarkdown scripts that are organized into analysis modules that generate curated reports. The modular framework is highly flexible and does not require complete reliance on a single analysis platform. Additionally, the self-contained reports generated by each module provide a comprehensive analysis summary and log of analytic parameters and scripts, thereby ensuring reproducible and shareable analysis workflows.

In tandem to scPipeline, we developed the scMiko R package that comprises a collection of functions for application-specific scRNA-seq analysis and generation of scPipeline analytic reports. We describe and validate novel scRNA-seq methods implemented in scMiko that facilitate multi-level cellular and functional annotation. Specifically, using eight reference scRNA-seq datasets (**Table 1**), we validate the co-dependency index (CDI) as a differential expression (DE) method that identifies binary differentially-expressed genes (bDEGs), propose a specificity-based resolution criterion to identify optimal cluster configurations, describe the Miko scoring pipeline for cell-type annotation, and introduce scale-free shared nearest neighbor network (SSN) analysis as a gene program discovery algorithm.

**Table 1.** Public scRNA-seq datasets used in the current study.

| Table 1. Public scRNA-seq datasets used in the current study. |  |  |  |  |  |  |
| --- | --- | --- | --- | --- | --- | --- |
| Dataset | Description | Species | Method | N |  | Analyses |
|  |  |  |  | Cells (% subset) | Types |  |
| Tabula Muris<sup>59</sup> | Pan-atlas | Mm | 10X | 100,000 (99%) | 100 | A, B, C |
| Tabula Sapiens<sup>60</sup> | Pan-atlas | Hs | 10X | 100,000 (21%) | 158 | A, B, C |
| Cao 2019<sup>3</sup> | Organogenesis | Mm | sci-RNAseq3 | 50,000 (100%) | 37 | A, B, C, E |
| Cao 2020<sup>57</sup> | Fetus | Hs | sci-RNAseq3 | 100,000 (26%) | 77 | A, B, C, E |
| Pijuan-Sala 2019<sup>24</sup> | Gastrulation | Mm | 10X | 100,000 (77%) | 38 | A, B, C, D, E |
| Tyser 2021<sup>16</sup> | Gastrulation | Hs | SMART-seq2 | 1,195 (100%) | 18 | A, B, C, E |
| La Manno 2021<sup>27</sup> | Developing brain | Mm | 10X | 100,000 (39%) | 16, 136 | A, B, C, E |
| Zeisel 2018<sup>58</sup> | Adolescent brain | Mm | 10X | 22,238 (100%) | 39 | A, B, C, E |
| Han 2022 (*in review*) | neural diff. | Mm | sci-RNAseq-3 | 26,117 (100%) | - | E |
| Ochocka 2021<sup>26</sup> | immune cells | Mm | 10X | 40,401 (100%) | - | E |

Analyses in which the data sets were used are indicated as *A*: DE methods, *B*: cluster resolutions, *C*: cell type gene sets, *D*: Miko scoring, *E*: gene program discovery.
Abbreviations: Hs; Homo sapiens (human), Mm; Mus musculus (mouse)

The scMiko R package (https://github.com/NMikolajewicz/scMiko) and scPipeline scripts (https://github.com/NMikolajewicz/scPipeline) are available on GitHub. Step-by-step tutorials and documentation are also provided at https://nmikolajewicz.github.io/scMiko/.

## Results

### 1. Overview of scPipeline modules

Here we introduce scPipeline, a modular collection of R markdown scripts that generate curated analytic reports for scRNA-seq analyses (**Fig 1**). For a given gene expression matrix, the ***QC and preprocessing module*** performs data filtering (based on mitochondrial content and gene recovery) and normalizes the count matrix using the scTransform algorithm implemented in Seurat^12^. The module outputs a Seurat object (for downstream analyses), and a corresponding standalone HTML report that summarizes the results^13^. In the case of multiple scRNA-seq datasets (e.g., experimental replicates, multiple studies and/or public datasets), we provide an ***integration module*** that leverages the canonical correlation analysis (CCA) and reciprocal principal component analysis (rPCA) approaches implemented in Seurat to facilitate data integration for downstream analyses^5^. Once data has been preprocessed, cells are clustered using the ***cluster optimization module***, where we introduce a novel specificity-based criterion for identifying the optimal resolution for Louvain community-based clustering. For each candidate cluster resolution, we also report DEGs identified using the Wilcox and CDI DE methods, for which we highlight specific and distinct applications in our current work. Once the optimal cluster configuration has been identified, the annotation modules facilitate cell type and cell state annotation using *a priori* cell-type markers, analysis of gene expression and associations, and unsupervised gene program discovery and functional annotation. Notably, the ***cell annotation module*** utilizes our novel gene set scoring method (i.e. the Miko score) to reliably annotate cell clusters using cell-type-specific markers. The Miko score is distinct from existing gene set scoring methods in that it adjusts for inherent variations in gene set size, thereby enabling direct comparison and ranking of gene set scores computed across gene sets of varying size. To facilitate gene expression exploration, we also developed a ***gene expression and association module*** which enables users to explore the expression pattern of query genes and predict gene function based on gene co-similarity profiles. Similarity profiles can be constructed using various methods, including Spearman correlation, rho proportionality, and CDI metrics^14^. These profiles are then functionally annotated to identify putative pathways correlated with the gene of interest. Finally, the ***gene program discovery module*** is used for gene program detection and transcriptomic network visualization. In addition to providing validated gene program discovery methods (e.g., ICA and NMF), we introduce the scale-free shared nearest neighbor network (SSN) method, which we demonstrate has superior recovery of known gene ontologies (GO) and enrichment of STRING-curated protein-protein interactions (PPI). Collectively, scPipeline offers a streamlined and reproducible workflow with user-friendly and intuitive reports and contributes to the current computational resources available for scRNA-seq. Importantly, its modular framework provides a foundation upon which future analysis modules can be developed to support additional scRNA-seq analyses.

**Figure 1.**
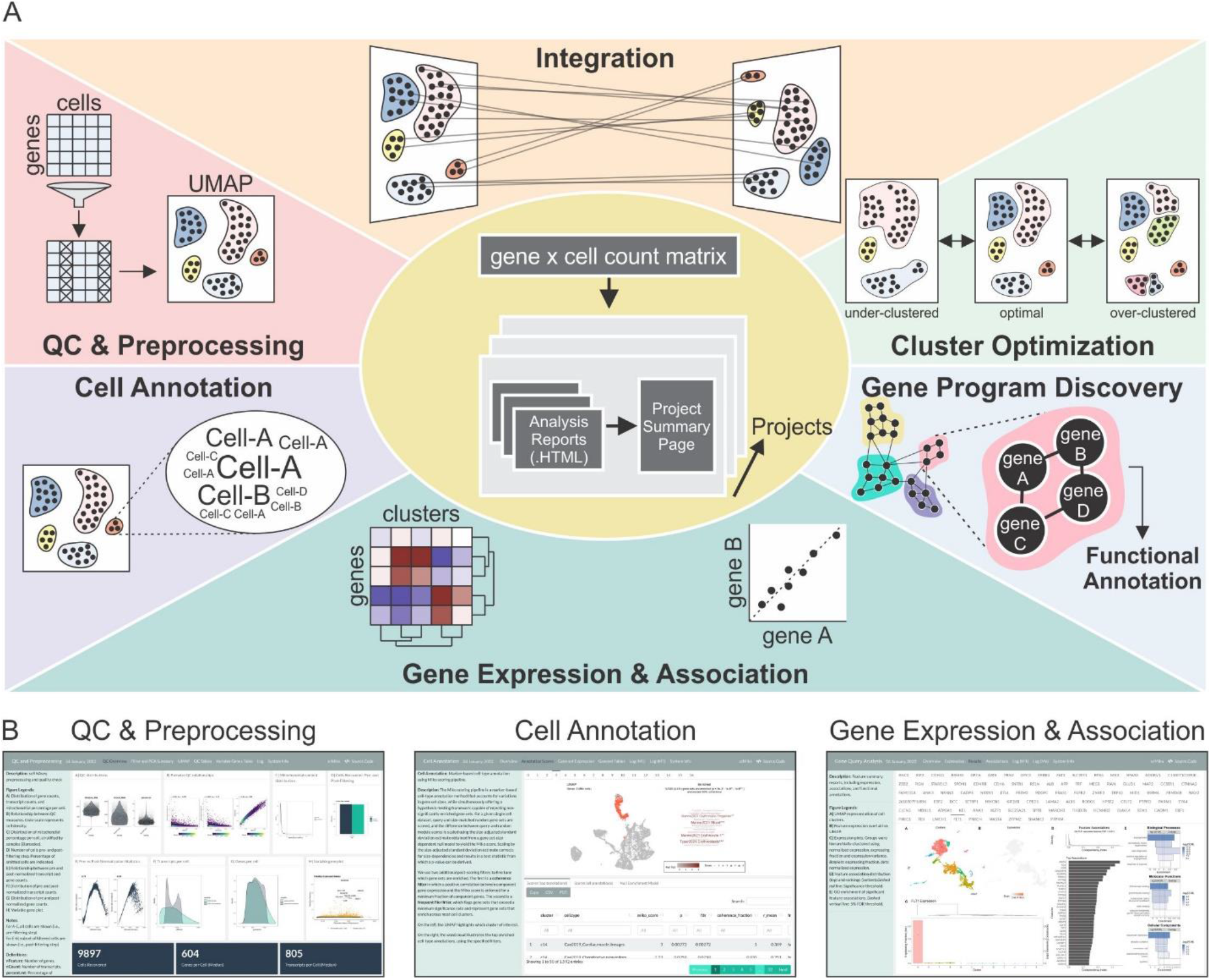
Schematic of scPipeline analysis modules. **(A)** scPipeline is a modular collection of Rmarkdown scripts that generate reports for scRNA-seq analyses. The modular framework permits flexible usage and facilitates *i*) QC & preprocessing, *ii*) integration, *iii*) cluster optimization, *iv*) cell annotation, *v*) gene expression and association analyses, and *vi*) gene program discovery. Each standalone .HTML report provides a comprehensive analysis summary that can be seamlessly shared without any dependencies. Alternatively, online repositories (e.g., GitHub) can be used to host .HTML reports for public dissemination. **(B)** Representative snapshots of scPipeline reports generated using the QC and preprocessing (*left*), cell annotation (*middle*), and gene expression and association (*right*) modules. More examples can be found [here](https://nmikolajewicz.github.io/scMiko/index.html).

### 2. Co-dependency index identifies cell-type specific markers

Robust identification of DEGs between cell populations is critical in scRNA-seq analyses. DEGs can be further subclassified into two different groups: graded DEGs (gDEG), in which genes are expressed in both populations, but to varying degrees; and binary DEGs (bDEG), in which genes are exclusively expressed in one population but not the other (**Fig 2A**). Popular scRNA-seq DE methods, such as the Wilcoxon method^15^, identify DEGs indiscriminately and require additional downstream filters to parse out bDEGs. Thus, a method tailored towards specifying bDEGs is needed.

**Figure 2.**
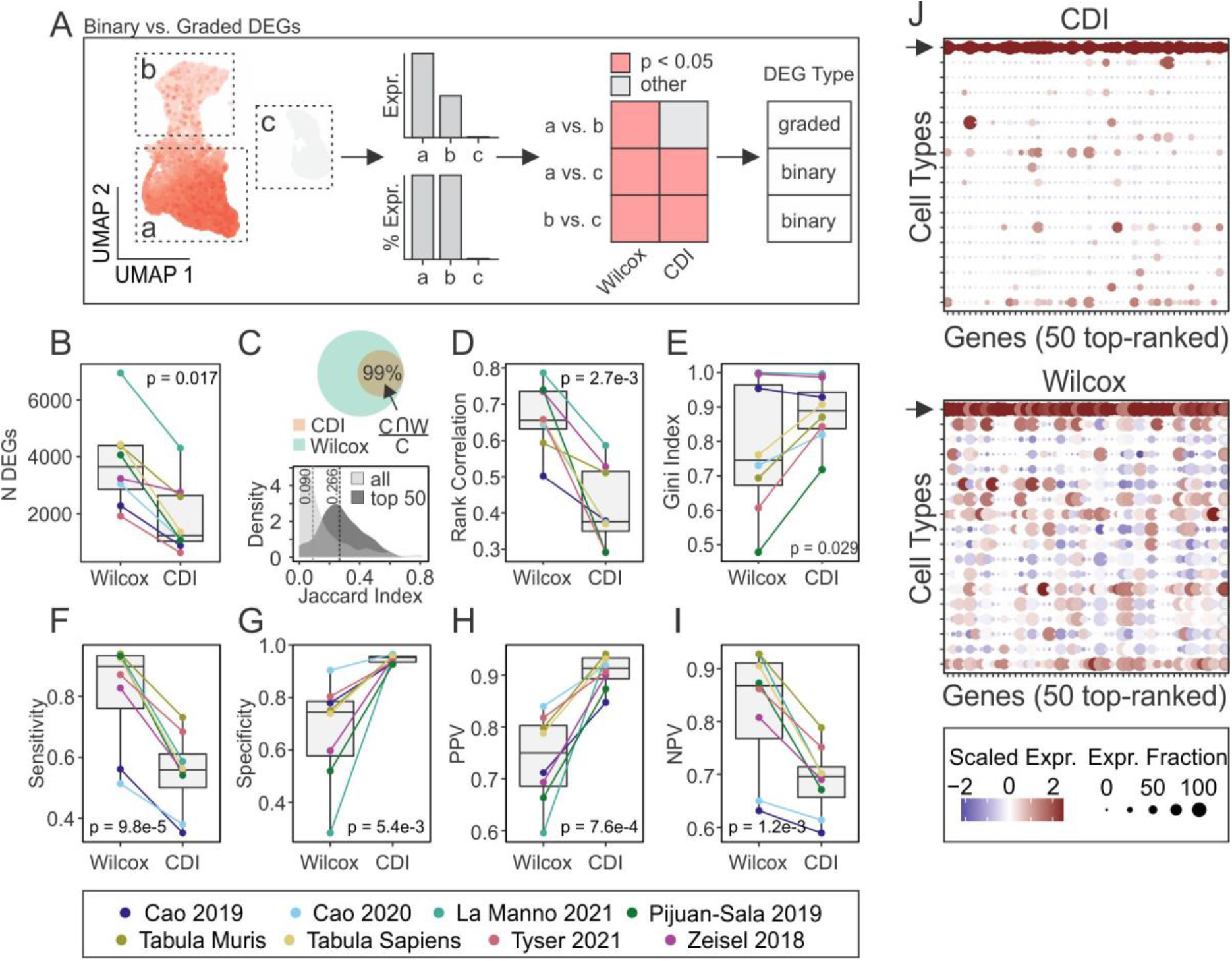
Co-dependency index identifies cell-type specific markers. **(A)** Schematic illustrating binary and graded DEGs in scRNA-seq analysis. (**B-I)** DEGs were identified by Wilcoxon and CDI methods across eight public scRNA-seq datasets and evaluated for number of significant DEGs (5% FDR, **B**), DEG overlap (**C**), rank correlation of average gene expression with -log10(p) values (**D**), Gini inequality index (**E**), sensitivity (**F**), specificity (**G**), positive predictive value (PPV, **H**), and negative predictive value (NPV, **I**). For **C**, the *top panel* shows overlap between DEGs identified by Wilcoxon and CDI, whereas the *bottom panel* shows the distribution of Jaccard similarities across all significant DEGs and top 50 DEGs. For **E-I**, the top 50 DEGs identified by each method were considered. **(J)** Representative dot plot of top 50 DEGs identified by CDI (*top*) and Wilcoxon (*bottom*) methods in yolk sac mesoderm cell population from Tyser 2021 scRNA-seq data (*arrows* indicate row corresponding to the yolk sac mesoderm population). For all comparisons, p values were determined by paired Wilcoxon ranked sum test. CDI; co-dependency index, DEGs; differentially-expressed genes, FDR; false discovery rate.

Here we propose using the CDI to identify cluster-specific bDEGs within scRNA-seq data. Using eight diverse public scRNA-seq datasets (**Table 1**), we identified significant DEGs using the CDI and Wilcoxon methods, and evaluated each method’s relative performance and behavior. The CDI method identified 66% fewer DEGs than the Wilcoxon method (1241 vs. 3653 genes, p = 0.017) (**Fig 2B**). These results reflect that the Wilcoxon method has the tendency to indiscriminately identify both gDEGs and bDEGs, whereas CDI selectively identifies bDEGs (**Fig 2C**, *top*). Among all the significant DEGs obtained by either method, the median Jaccard similarity was 0.09; however, when only the top 50 DEGs [ranked by -log10(p)] were considered, the Jaccard similarity increased to 0.266, suggesting a bias towards bDEGs among top DEGs identified by Wilcoxon (**Fig 2C**, *bottom*). Consistent with prior reports, the Wilcoxon method was systematically biased towards calling highly-expressed genes differentially-expressed. While this bias was present for the CDI method, it was significantly lower in contrast to the Wilcoxon method (**Fig 2D**, p = 2.7e-3), and in the range of the best performing methods evaluated previously^15^. Finally, we evaluated the cluster-discriminating accuracy of the top 50 genes identified by each method (**Fig 2E-I**). While the Wilcoxon method identified genes with higher cluster-discriminating sensitivity (0.90 vs. 0.56, p = 9.8e-5; **Fig 2F**) and negative predictive value (NPV; 0.87 vs. 0.70, p = 1.2e-3; **Fig 2I**), the CDI method had superior specificity (0.95 vs. 0.75, p = 5.4e-3; **Fig 2E, G**) and positive predictive value (PPV; 0.91 vs. 0.75, p = 7.6e-4, **Fig 2H**). As an illustrative example, we evaluated the top 50 DEGs in yolk-sac mesoderm^16^, where we observed a higher degree of specificity among the top markers identified by the CDI method (**Fig 2J**). Together, these analyses establish the CDI method as an approach to specifically identifying bDEGs.

### 3. Marker specificity-based criterion for identifying optimal cluster resolutions

scRNA-seq-based cell type identification relies on unsupervised clustering methods; however, resulting cell clusters can vary drastically depending on what resolution is used to perform clustering. Many approaches have been proposed to guide the selection of the optimal resolution, including silhouette index^17^ and resampling-based methods (e.g., chooseR^18^ and MultiK^19^). However, these methods are motivated by theoretical rather than biological criterion. Having demonstrated that the CDI method yields cluster-specific markers (**Fig 2**), we propose to define cell-types at a clustering resolution that maximizes the specificity of markers obtained in each cluster. We proceed by first clustering over a range of candidate resolutions, and the top specific marker in each cluster at each resolution is identified using the CDI method (**Fig 3A**, *step 1*). Subsequently, specificity curves are generated for each resolution and used to obtain aggregate specificity metrics. The resolution at which maximal specificity is observed is taken as the optimal resolution, *S*_*peak*_ (**Fig 3A**, *step 2*). However, acknowledging that there exist multiple levels of resolution that are biologically relevant (e.g. cell types vs. cell subtypes)^19^, we observed that the specificity curves in many datasets exhibited “elbows”, which we hypothesize represent additional biologically relevant clustering configurations, and we termed these *S*_*elbow*1_ and *S*_*elbow*2_.

**Figure 3.**
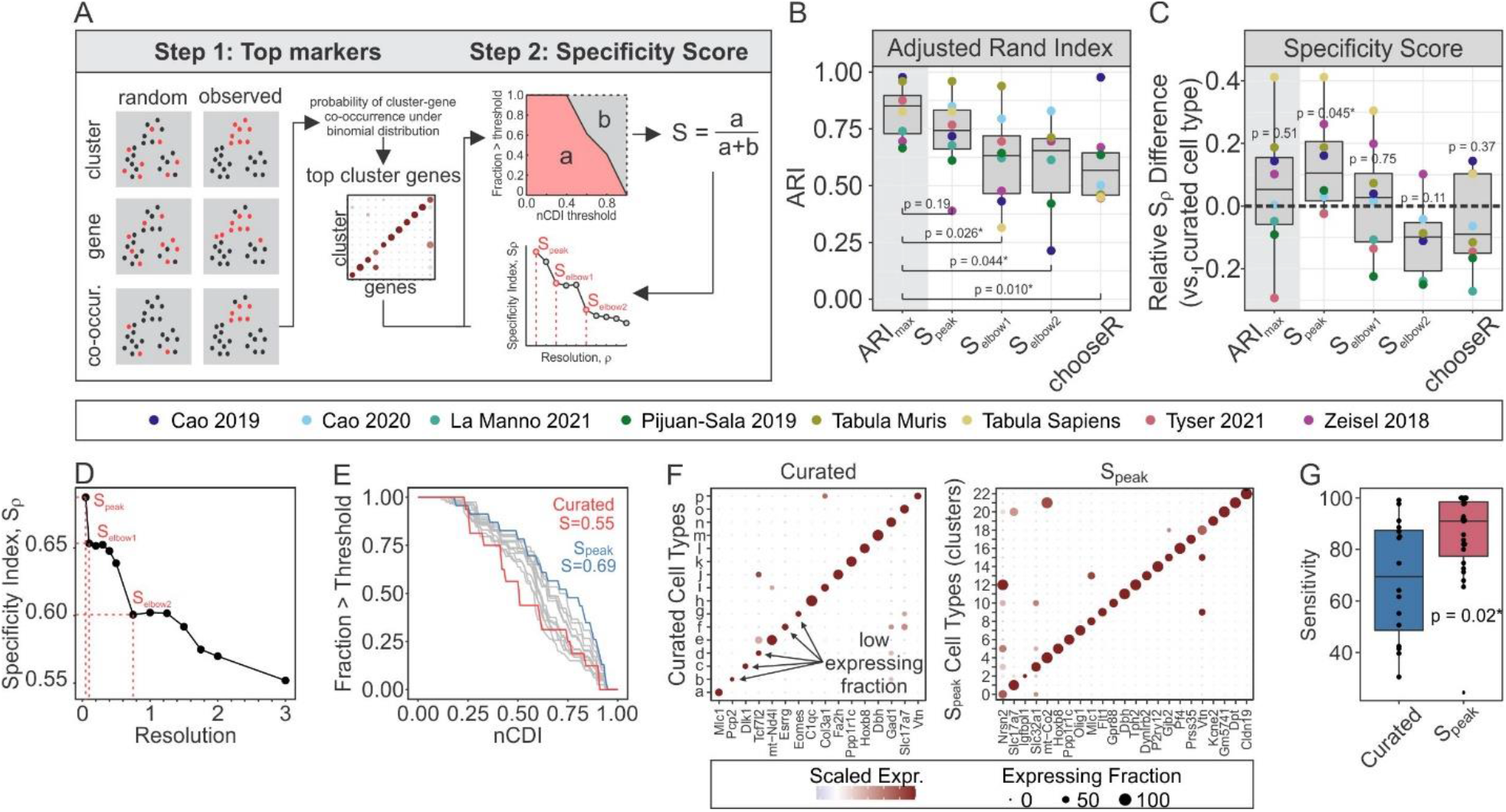
Identification of optimal clustering resolution using a specificity-based criterion. **(A)** Schematic of specificity-based resolution-selection criteria. **(B, C)** Adjusted Rand index (**B**) and specificity index differences (**C**) between ground truth (author-curated) clusters and observed Louvain clusters, using resolutions determined by different optimization criteria [specificity criteria (*S*_*peak*_, *S*_*elbow*1_, *S*_*elbow*2_) and *chooseR*^18^]. *ARI*_*max*_ represents resolution at which maximal ARI was achieved, after considering all candidate resolutions (0.5 to 3). For **B**, significance compared to *ARI*_*max*_ was determined by paired Wilcoxon test. For **C**, significance compared to zero (i.e., ground truth) was determined by one-sample Wilcoxon test. **(D-G)** Optimal clustering resolution for Tyser 2021 human gastrulation scRNA-seq data^16^. **(D)** Relationship between resolution and specificity indices, and identification of *S*_*peak*_, *S*_*elbow*1_ and *S*_*elbow*2_. **(E)** Specificity-curves. *Grey curves*: all candidate resolutions evaluated (0.5 to 3), *blue curve*: S_peak_, *red curve*: ground truth (curated) clusters. **(F)** Dot plots of top markers for curated (*left*) and *S*_*peak*_ (*right*) clusters. **(G)** Comparison of cluster sensitivity (expressing fraction) of each top marker obtained for curated and *S*_*peak*_ clusters. Significance determined by unpaired Wilcoxon test

To evaluate the performance of our specificity-based resolution selection criteria (*S*_*peak*_, *S*_*elbow*1_, and *S*_*elbow*2_), we used eight public scRNA-seq datasets, and adopted author-curated cell-types as “ground-truth” clusters. We showed that our specificity-based criteria favor clustering configurations that align with manually curated cluster labels, as indicated by the lack of significant difference between the adjusted Rand index (ARI; i.e., a measure of classification consistency) obtained at *S*_*peak*_ and *ARI*_*max*_ resolutions (**Fig 3B**). By comparison, *chooseR* (a resampling-based resolution selection criteria), *S*_*elbow*1_ and *S*_*elbow*2_ yielded clusters with significantly lower ARI, suggesting that these cluster configurations represent cell subtypes, whereas clusters obtained at the *S*_*peak*_ resolution represent well-defined cell type clusters (**Fig 3B**). In support of this, *S*_*peak*_ clusters were associated with significantly more specific markers (i.e., top markers were more specific) than “ground truth” clusters (p = 0.045), whereas there was no significant difference observed for the other cluster configurations compared to “ground truth” clusters. As a representative example, we applied our specificity-based resolution selection approach to the human gastrulation scRNA-seq data published by Tyser and colleagues (2021)^16^ (**Fig 3D**). Compared to curated clusters, *S*_*peak*_ clusters were associated with a higher specificity index (0.69 vs. 0.56) (**Fig 3E**) which was verified by visual inspection (**Fig 3F**), and further, it was demonstrated that the top markers associated with *S*_*peak*_ clusters were significantly more sensitive (i.e., high expression fraction; p = 0.02) than those obtained in “ground truth” clusters (**Fig 3G**). Our results demonstrate that a specificity-based resolution selection criterion reliably identifies cluster configurations that reflect biologically relevant cell types.

### 4. Marker-based cluster annotation with Miko score

Transcriptome-wide expression profiling has led to the generation and availability of gene sets for cell-type identification. Nonetheless, the external validity of these genes sets is remarkably inconsistent, largely stemming from the fact that many gene sets are derived using one-versus-all DE methods on genetic backgrounds that lack population-level phenotypic diversity. While elucidating the exact conditions under which a gene set reliably identifies a given cell type is beyond the scope of the current study, we argue that cell-type specific gene sets obtained using one-versus-all DE methods are most valid when derived from diverse cell atlases. To complement our marker-based cluster annotation efforts, we performed DE analysis on the eight public scRNA-seq datasets presented in **Table 1**, each comprising highly diverse cell types. Together with cell type markers reported in Zhao 2019^20^ and the PanglaoDB^21^, we provide a catalog of cell type markers comprising 1043 (redundant) cell type-specific marker sets spanning 11748 unique genes. Representating the cell-type marker catalog as a bipartite network revealed major cell type hubs including epithelial, mesenchymal, endothelial, and lymphoid/hematopoietic cell types, in addition to tissue-specific cell ontologies like cardiac, neural, and glial cells (**Fig 4A**).

**Figure 4.**
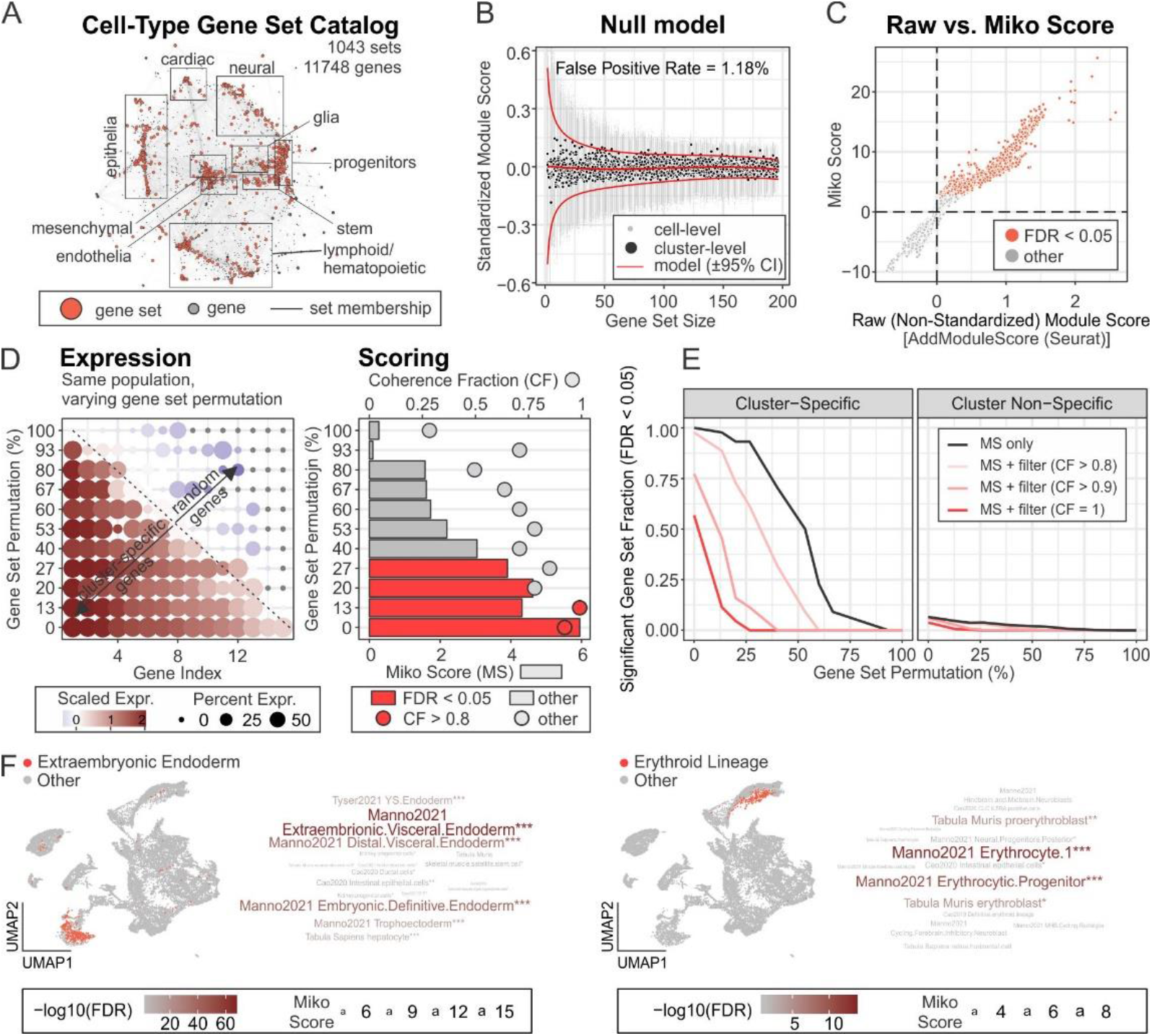
Cell-type annotation. **(A)** Cell-type-specific gene set catalog represented as bipartite network. Edges between gene sets (*red nodes*) and genes (*grey nodes*) represent gene set membership. Major cell ontologies are annotated, and the corresponding gene sets can be accessed using the scMiko R package. **(B)** Representative null model relating gene set size and standardized module scores (SMS; for random gene sets). *Red curves*: predicted mean SMS ± 95% CI; *black points*: observed cluster-level mean SMS; *grey points*: observed cell-level SMS. **(C)** Relationship between cluster-level non-standardized module score (AddModuleScore, Seurat R package) and Miko score. Clusters with significant module activity (FDR < 0.05) are indicated. **(D-E)** Evaluation of Miko score performance. **(D)** Representative gene sets with varying rates of permutation (i.e., substitution of cluster-specific gene with random gene; *left*) and corresponding Miko scores (bar plot, *right*) with coherent fractions (dot plot, *right*). **(E)** Relationship between degree of gene set permutation and fraction of cluster- and cluster-non-specific gene sets with significant (FDR < 0.05) module activity. Coherent fraction (CF) filters were included to demonstrate capacity to titrate score sensitivities and specificities. **(F)** Representative example of Miko score applied to murine gastrulation data using cell-type gene set catalog (**A**). UMAPs illustrate cell population with curated cell-types of interest (Extraembryonic endoderm; *left*, erythroid lineage; *right*), and word clouds represent top cell types predicted by the Miko scoring algorithm.

Many marker-based cell annotation methods have been described^22,23^; however, one limitation of these methods is a lack of consideration for gene set size. As the number of genes in a gene set increases, pooled signature scores become less sensitive to the influence of highly expressed individual genes. This gene set size dependency leads to a bias, such that scores obtained from smaller gene sets tend to have more spurious enrichments than those obtained from larger gene sets (**Fig 4B**), precluding unbiased comparison of signature scores obtained over a range of unevenly sized gene sets. Motivated by this limitation, we introduce the Miko score, a cell cluster scoring method that accounts for variations in gene set sizes. The Miko score also provides a hypothesis-testing framework capable of rejecting non-significantly enriched gene sets (**Fig 4**). For a given single-cell dataset, query and size-matched random gene sets are scored using a standardized implementation of AddModuleScore(…), and the difference between query and random module scores is scaled using the size-adjusted standard deviation estimate obtained from a gene set size-dependent null model (**Fig 4B**) to yield the Miko score (**Fig 4C**). The standardized implementation of AddModuleScore(…) accounts for cell-to-cell variation in gene expression, while scaling by the size-adjusted standard deviation estimate adjusts for size-related dependencies and results in a test statistic from which a p-value can be derived.

The performance of Miko score-based cell annotation was evaluated using cell-type-specific gene sets derived for each cell type in the mouse gastrulation dataset reported by Pijuan-Sala and colleagues^24^. To assess the robustness of the Miko score and account for inaccuracies in gene set definitions, each set was permuted to varying extents, such that a subset of cell-type specific markers in each gene set were replaced with an equal number of randomly sampled genes (**Fig 4D**). Using non-permuted gene sets, the Miko score-based enrichments were 100% sensitive and 94% specific for cluster-specific gene sets (**Fig 4E**). When 25% of genes were permuted, we observed 93% sensitivity and 96% specificity. However, at higher permutation rates, we observed a significant decline in sensitivity such that at 50% permutation there was 54% sensitivity and 98% specificity. We also found that filtering enrichments using a coherence criterion resulted in marginally improved specificity at the cost of sensitivity (**Fig 4E**). As an illustrative example, we calculated Miko scores using our cell-type marker catalog (**Fig 4A**; Pijuan-Sala-derived markers were omitted from the catalog) and demonstrated that author-curated endoderm and erythroid populations were accurately annotated using our Miko score pipeline (**Fig 4F**). Collectively, our analyses establish the Miko score as a marker-based scoring algorithm that is robust to gene set inaccuracies and capable of facilitating unbiased comparison across a large collection of unevenly sized gene sets.

### 5. Gene program discovery using scale-free topology shared nearest network analysis

Unsupervised gene program discovery offers a complementary approach to annotating cell clusters in scRNA-seq, which aim to group genes based on co-expression similarity profiles. Here we introduce the scale-free topology shared nearest network (SSN) method to identify gene expression programs (**Fig 5A**). In brief, the gene expression matrix is dimensionally reduced using principal component analysis (PCA). Each gene’s K-nearest neighbors (KNN) is then determined by Euclidean distance in PCA space. The resulting KNN graph is used to derive a shared nearest neighbor (SNN) graph by calculating the neighborhood overlap between each gene using the Jaccard similarity index. Adopting the framework from weighted gene correlation network analysis (WGCNA)^25^, an adjacency matrix that conforms to a scale-free topology is then constructed by raising the SNN graph to an optimized soft-thresholding power, which effectively accentuates the modularity of the network (**Fig 5B**). The resulting adjacency matrix is used to construct the network UMAP embedding and to cluster genes into programs (or modules) by Louvain community detection. To reduce noise, genes with low connectivity (i.e., low network degree) are pruned so that only hub-like genes are retained for downstream annotation and analysis.

**Figure 5.**
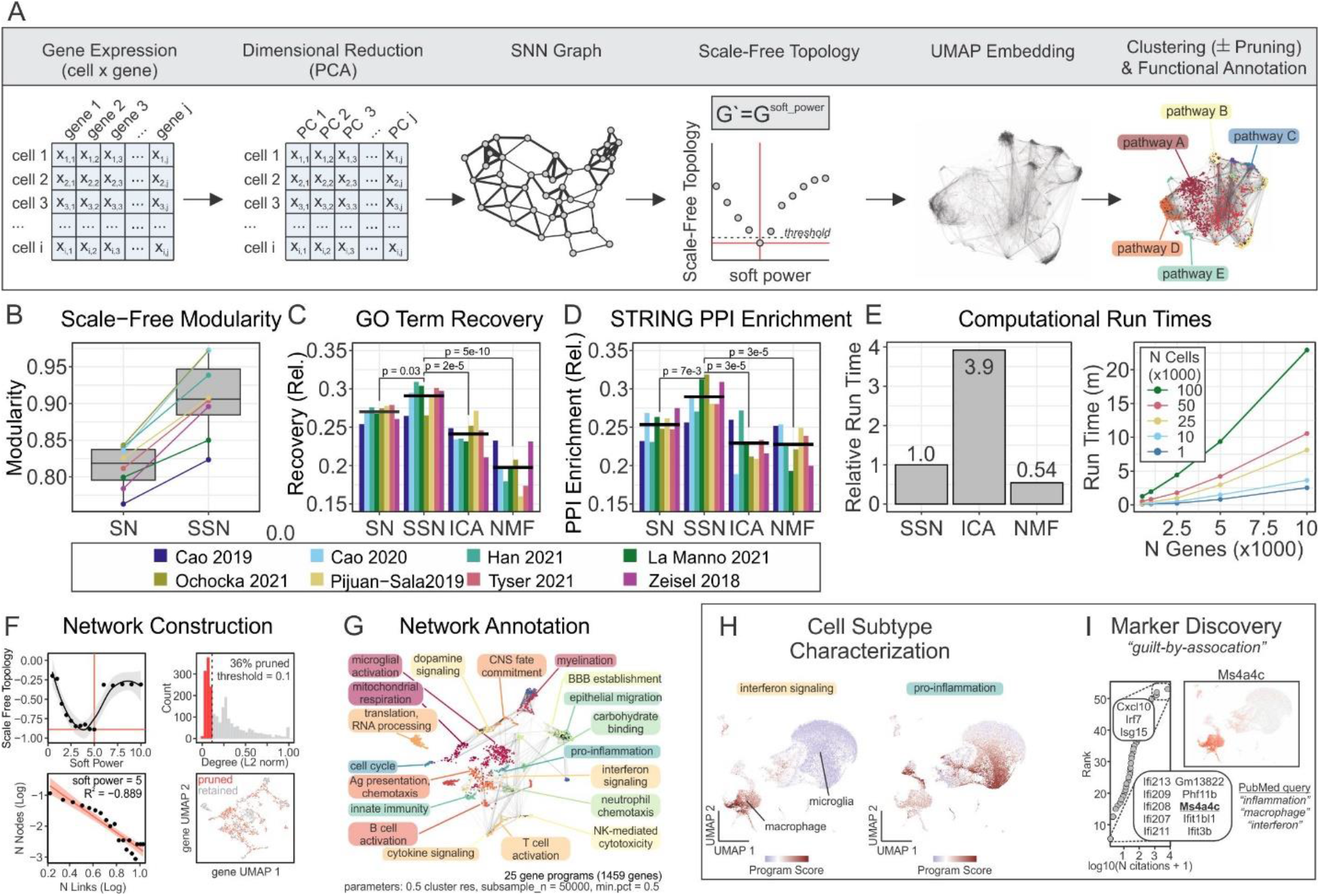
Gene program discovery using scale-free shared nearest neighbor network (SSN) analysis. **(A)** Schematic illustrating network construction and annotation. **(B)** Network modularity with (SSN) and without (SN) scale-free topology enforcement. **(C-E)** Comparison of GO term recovery (**C**), STRING PPI enrichment (**D**) and computational run time (**E**) across different gene program discovery methods. ICA; independent component analysis, NMF; non-negative matrix factorization, SN; shared nearest neighbor network, SSN; scale-free shared nearest neighbor network. (**F-I**) Representative transcriptional network construction, annotation and applications using Ochocka 2021 scRNA-seq data^26^. **(F)** Optimal soft power required for scale-free topology (*left column*; threshold = -0.9) and pruning of genes with low network connectivity (*right column*; threshold = 0.1). **(G)** Functional annotation of gene programs. GO term enrichment was performed using hypergeometric overrepresentation analysis. **(H)** Activity of “interferon signaling” and “pro-inflammation” programs overlaid on cell UMAP. Macrophage and microglial subpopulations can be subtyped by program activity status. **(I)** Novel marker discovery and functional prediction using guilt-by-association. Genes belonging to “interferon signaling” program were cross-referenced with PubMed articles queried using “inflammation”, “macrophage” and “interferon” search strings to identify novel candidate genes (e.g., Ms4a4c) implicated in interferon signaling. Ms4a4c expression was visualized on a UMAP to verify that expression is coherent with gene program activity.

Compared to independent component analysis (ICA) and non-negative matrix factorization (NMF), SSN gene programs had significantly superior GO term recovery and STRING PPI enrichment (**Fig 5C, D**). The importance of enforcing a scale-free topology was evident in the comparison between SN (shared nearest neighbor network *without* scale-free topology) and SSN (shared nearest neighbor network *with* scale-free topology) (**Fig 5C, D**). On average, the relative computational runtimes were 0.54, 1, and 3.9 for NMF, SSN, and ICA methods, respectively, thereby establishing NMF as the fastest algorithm, but only by a small margin over SSN which significantly outperformed ICA (**Fig 5E**).

We demonstrated the use of SSN gene program discovery and network visualization with two case examples (**Fig 5F-L**). In the first case, we constructed an SSN network using scRNA-seq data of the murine immune compartment in brains engrafted with the syngeneic GL261 GBM cell line^26^ (**Fig 5F**). Functional annotation of each gene program revealed a diverse transcriptomic landscape (**Fig 5G**), including interferon signaling and pro-inflammatory programs that were highly active in monocyte/macrophage and microglial sub-populations, respectively (**Fig 5H**). In addition to facilitating further cellular characterization, functionally annotated gene programs offer opportunities to predict the function of previously uncharacterized genes using a “guilt-by-association” approach. For example, cross-referencing genes belonging to the interferon-signaling gene program in the SSN graph with PubMed-indexed publications, we find the gene *Ms4a4c* had not been previously associated with “inflammation”, “macrophage” or “interferon”. We predict that *Ms4a4c*, a previously uncharacterized gene, may have a role in the inflammatory process (**Fig 5I**). In our second example, we demonstrate how SSN gene program discovery can identify and facilitate the refinement of robust gene signatures (**Fig S2**). Briefly, we constructed a SSN network from scRNA-seq data derived from a murine developing brain^27^ (**Fig S2A-B**) and show that the expression of each gene belonging to the angiogenesis program is positively correlated with the aggregate gene program score when examined in the developing murine brain data from which the signature was derived (**Fig S2C**, *left*). Notably, in two other independent datasets (murine and human gastrulation), only a subset (albeit majority) of genes were positively correlated with the program score (**Fig S2C**, *middle, right*). By taking the 3-way intersection of coherent genes across these three relevant datasets, we find a 64-gene signature (**Fig S2D**) that was specifically enriched among the hematogenic endothelial populations in all three scRNA-seq datasets (**Fig S2E**). Further supporting the validity of this gene signature refinement strategy, we previously applied this approach in the context of glioblastoma, where we derived robust prognostic signature panels that validated across multiple independent patient cohorts.^28^

## Discussion

### Overview

We have described a pair of computational resources, scMiko (R package) and scPipeline (dashboard analysis reports), and propose new methods to facilitate multiple levels of cluster annotation in scRNA-seq data. Our computational tools follow established scRNA-seq analytic practices, and offer modular workflows that enable data preprocessing, normalization, integration, clustering, annotation, gene program discovery and gene association analyses. Among the novel methods presented in this work, we validated the CDI as a DE method that identifies binary DEGs. Given the inherent specificity of bDEGs, we then adopted the CDI algorithm to derive a specificity-based resolution selection criterion for determining optimal clustering configurations and benchmarked the performance of this approach against ground truth annotations. Upon identifying the optimal cluster resolution(s), we demonstrate how to annotate clusters using our Miko Scoring pipeline, which facilitates unbiased scoring of a diverse set of variable-sized cell-type-specific gene sets and accepts or rejects candidate annotations using a hypothesis-testing framework. Finally, we describe scale-free shared nearest neighbor network (SSN) analysis as an approach to identify and functionally annotate gene sets in an unsupervised manner, providing an additional layer of functional characterization of scRNA-seq data.

### Annotation methods

The annotation methods presented here, namely finding bDEGs with CDI, cell-type annotation with Miko Scoring, and gene program discovery and functional annotation with SSN analysis, all complement and expand the extensive list of analytic methods for scRNA-seq^29,30^. It has become evident from systematic benchmarking efforts that no single method is enough to probe single cell datasets in-depth, and that several methods offer unique advantages with regards to biological accuracy, interpretability, computational complexity, visualization, or accessibility^14,15,31^.

#### Cluster optimization

Reliable annotation begins with identifying the optimal clustering configuration. Although there are many ways to cluster single-cell data, including K-means (SAIC^32^, RaceID3^33^), hierarchical (CIDR^34^, BackSPIN^35^, SINCERA^36^) and density-based (Monocle2^37^, GiniClust^38^) clustering, we used the community-detection based Louvain approach implemented in Seurat due to its low run time and high performance index^39,40^ and focused on optimizing the resolution that controls the number of resolved clusters. If cells are clustered at an inappropriately low resolution (i.e. under-clustered), there is a risk of amalgamating distinct cell types into single populations, resulting in a loss of resolution in cellular identity. In contrast, if the resolution is too high (i.e. over-clustered), multiple near-identical cellular lineages emerge and obscure the true complexity of the dataset. At the same time, it is recognized that clustering configurations at multiple different resolutions may be biologically relevant, and reflect different layers of cellular identities, such as cell types at lower resolutions (e.g., macrophage), and cellular sub-types (M1 vs. M2 polarized macrophage) at higher resolutions^19^. There are different selection criteria for identifying the optimal resolution(s), including the silhouette index^17^ and resampling-based methods (e.g., chooseR^18^, MultiK^19^); however, these methods are motivated by theoretical rather than biological criterion. The specificity-based resolution selection criterion described in our current work identifies cluster configurations coinciding with maximal marker specificity. This is a desirable property for downstream applications that require individual biomarkers to resolve cell types, such as flow cytometry or imaging. Additionally, when evaluated over multiple candidate resolutions, more than one biologically relevant resolution is often identified, manifesting as “elbows” on the specificity-resolution curve (akin to the elbow method used for selecting the number of principal components on a Scree plot). We benchmarked the performance of our specificity-based criterion against author-curated “ground truth” annotations and demonstrated that a specificity-based criterion outperforms the resampling-based approach used in chooseR. We note that a limitation of our method relates to the stability and reproducibility of clusters, especially in single-replicate data sets. Artifact genes (i.e., genes that are highly expressed exclusively in a small subset of cells belonging to a single experimental replicate) have been shown to produce distinct cellular clusters and in the absence of experimental replicates, and it is difficult to determine whether these clusters represent technical artifacts or real biology^41^. While this can be addressed through profiling multiple experimental replicates^41^, it may also be circumvented by expanding our specificity-based criterion to consider the top 5-10 markers, rather than the top single cluster-specific marker, at each resolution. Finally, although we evaluated our specificity-based criterion using the Louvain clustering approach, the criterion may be applied to any clustering method that requires optimization of the number of resolved clusters (e.g., K-means). We expect that our specificity-based criterion will complement existing optimization methods to find meaningful cluster configurations.

#### CDI DE method

The CDI DE method offers an approach to identifying bDEGs, which have applications distinct from gDEGs. Whereas gDEGs are useful for identifying differences that occur on a spectrum (e.g., neural development), bDEGs have greater utility in identifying cell-type-specific markers (e.g., FACS sorting of CD34^+^ for hematopoietic stem cells), diagnostic biomarkers, disease targets (e.g., CART-cell therapy), and artifact genes in scRNA-seq datasets^41^. A known limitation of existing DE methods for scRNA-seq is the failure to account for variation in biological replicates, and the CDI approach is no exception^15^. Nonetheless, we expect that with appropriate biological replicates and external validation, the CDI DE method will contribute to the identification of specific biomarkers.

#### Cell annotation

The Miko scoring cell-type annotation workflow described in this work supplements the existing repertoire of marker-based annotation algorithms including scCatch^42^, SCSA^43^, SCINA^44^, and CellAssign^45^. The hypothesis testing framework implemented in the Miko scoring pipeline enables the rejection of unlikely cell-type annotations, a property that is shared by SCINA and CellAssign. However, unlike its predecessors, Miko scoring explicitly corrects for gene set size biases, thereby enabling unbiased comparison of scores over a large collection of various sized gene sets. This property enables prioritization of the most likely annotation if multiple marker sets are significantly scored for a given population. Coupled with our word cloud-based visualizations introduced in scMiko and scPipeline, candidate cell-type annotations can be easily inspected and interpreted.

#### Cell-type marker database

To facilitate marker-based annotation of cell types, several reference databases are available including CellMatch^42^, CellMarker^20^, PanglaoDB^21^, CancerSEA^46^, and MSigDB (collection 8)^47^. We contribute to these resources by deriving marker sets from diverse single-cell atlases (**Table 1**), and through network-based visualization we demonstrate the hierarchical complexity of cell ontology (**Figure 4A**). While the network organization was generally coherent with the cell-type annotations assigned to the marker sets, an inspection of select local neighborhoods in our cell-type marker network revealed occasional co-similarities between marker sets from heterogeneous cell types, reflecting either inaccuracies in marker curation or similarities in cellular processes across dissimilar cell types. Based on these observations, we emphasize that marker-based annotations are only as good as the cell-type prescribed to the original dataset. Thus, integrating a large collection of marker sets from multiple independent sources to achieve consensus annotations, or alternatively, using a robustly validated collection of marker sets can attain optimal results.

#### Gene program discovery

The SSN method for gene program discovery was inspired by the established shared-nearest neighbor (SNN) framework used in single-cell analyses to reliably identify cell-to-cell distances in a sparse dataset, as well as the scale-free topology transformation used under the assumption that the frequency distribution of gene association in a transcriptomic network follows the power law^25,48,49^. A UMAP-embedded network, based on a SNN graph akin to that used in our SSN procedure, has previously resolved gene modules corresponding to protein complexes and pathways, with Euclidean distances in UMAP space out-performing correlation and PCA distances in predicting protein-protein interactions ^50^. Consistent with these findings, we demonstrated that gene programs identified by SSN yielded superior GO term recovery and enrichment of STRING PPIs compared to ICA and NMF methods, and that the scale-free topology transform was critical in driving this improvement in performance. Taken together, the SSN gene program discovery method is robust to data sparsity, has a high performance index, offers a network-based visualization, and has run-times that scale well for larger datasets.

### Concluding Remarks

Future plans for scPipeline and scMiko involve continual review and improvement of existing workflows, and development of additional analysis modules that facilitate complementary analyses such as characterization of ligand-receptor interactions^51,52^, regulon-based transcription factor inference^53^, trajectory analyses^3,54,55^ and differential-abundance analyses^56^. As innovative approaches to interrogate single cell data are proposed by us and others, we will adopt and implement these for all users to benefit.

## Methods

### Software

Figure preparation: CorelDRAW x8 (Corel); Bioinformatic analyses: R v 4.0.3 (R Foundation for Statistical Computing).

The scMiko R package and scPipeline are freely available and documentation and tutorial vignettes can be found here: https://nmikolajewicz.github.io/scMiko/.

### Data sources

scRNA-seq data from Ochocka et al. (2021) was obtained from Gene Expression Omnibus (GEO; accession number GSE136001)^26^; Cao et al. (2019) from GEO (accession number GSE119945)^3^; Cao et al. 2020 from GEO (accession number GSE156793)^57^; Zeisel et al. (2018) from http://mousebrain.org/downloads.html^58^; La Manno et al. (2021) from http://mousebrain.org/downloads.html^27^; Tabula Muris from [FigShare](https://figshare.com/articles/dataset/Robject_files_for_tissues_processed_by_Seurat/5821263)^59^; Tabula Sapiens from [FigShare](https://figshare.com/articles/dataset/Tabula_Sapiens_release_1_0/14267219)^60^; Pijuan-Sala (2019) from the [MouseGastrulationData](https://github.com/MarioniLab/MouseGastrulationData) R Package^24^; and Tyser et al. (2021) from http://www.human-gastrula.net/^16^.

### Data preprocessing

scRNA-seq data sets were normalized, scaled, dimensionally reduced and visualized on a UMAP using the *Seurat* (v 4.0.4) workflow^4-7^. In brief, count matrices were loaded into a Seurat object and normalized using *NormalizeData*(…, normalization.method = ‘LogNormalize’, scale.factor = 10000). Variable features were identified using *FindVariableFeatures* (…, selection.method = ‘mvp’, mean.cutoff = c(0.1,8), dispersion.cutoff = c(1,Inf)) and then data were scaled using *ScaleData*(…). Principal component analysis, and UMAP embedding was performed using *RunPCA*(…) and *RunUMAP*(…, dims = 1:30), respectively. Metadata from original publications were used to annotate cell types.

### Statistical analyses

All pairwise comparisons were performed using the signed Wilcoxon rank sum test, and p values were adjusted for multiple comparisons using the Benjamini-Hochberg procedure, as indicated. In cases where methods were compared across a common set of data, paired Wilcoxon tests were performed.

### Differential expression analysis

Differential expression analyses were performed using Wilcoxon rank sum (Wilcox) and codependency index (CDI)^61,62^. The Wilcox method was implemented using the *wilcoxauc* function (*Presto* R package, v 1.0.0)^63^. Alternatively, the CDI was adopted to calculate the probability of cluster and gene co-occurrence under a binomial distribution. For a given gene *g* and cluster *k*, the joint probability of observed non-zero *g* expression in *k* is formulated as:

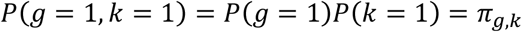

The probability of observing a test statistic more extreme under the null hypothesis that gene *g* and cluster *k* are independent is then:

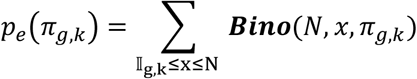

where ***Bino***(*N, x, π*_*g,k*_) represents the probability of observed *x* successes in *N* trials if the probability of success is *π*_*g,k*_, and 𝕀_g,k_ is the number of cells in which *g* and *k* are coincident. CDI is then defined as:

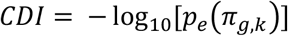

We further normalized the CDI score using the CDI score corresponding to the probability of observed a perfect co-dependency for cluster *k*:

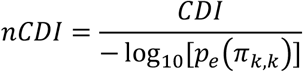

where *π*_*k,k*_ = *P*(*c*_*k*_ = 1, *c*_*k*_ = 1), under the assumption of independence. Possible values of *nCDI* range between [0,1], such that *nCDI* = 1 represents perfect co-dependence between a gene and cluster, and *nCDI* = 0 represents no co-dependence but is not equivalent to mutual exclusivity which has been formulated elsewhere^62^.

The CDI, by definition, only computes genes that are “up-regulated” relative to the comparison group, so to ensure fair comparison to the Wilcox method, only gene subsets that had a positive log fold change (LFC) were considered in Wilcox vs. CDI comparative analyses. Differentially expressed genes (DEGs) were deemed significant at a 5% false discovery rate (FDR). The top 50 DEGs identified by each method were subsequently characterized using sensitivity, specificity, positive predictive value (PPV) and negative predictive value (NPV):

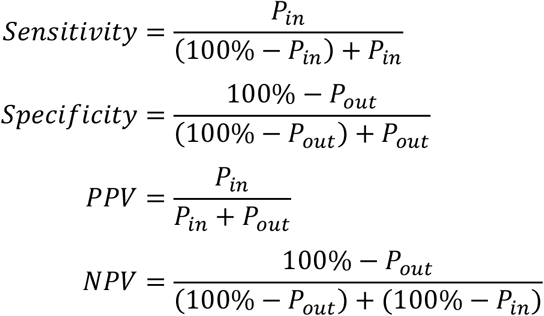

where *P*_*in*_ and *P*_*out*_ represent the expressing percentage of cells within and outside a cluster, respectively. We also computed the Gini inequality index as a complementary surrogate for gene specificity:

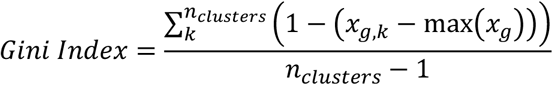

where *x*_*g,k*_ is the average expression of gene *g* for cluster *k*, and *n*_*clusters*_ is the number of unique clusters.

### Cluster optimization

To identify the optimal cluster resolution, we first clustered samples over a range of candidate resolutions (0.05 to 3) using *FindClusters*(…, algorithm = 1) in Seurat. At each resolution *ρ*, the top cluster-specific marker for each cluster was identified using CDI-based DE analysis. Subsequently, specificity curves were generated by plotting the proportion of clusters that exceed a threshold nCDI score, for nCDI ranging [0,1]. The area under this curve (AUC) represents the aggregate specificity index *S*_*ρ*_ and possible values range between [0,1], with a score of 1 representing the ideal cluster configuration in which each cluster has at least one marker satisfying nCDI = 1. Aggregate specificity indices were graphed over the range of candidate resolutions, and resolutions at which a peak and subsequent elbow(s) were manually observed were taken as optimal clustering resolutions for downstream analyses. Cluster resolutions were also identified using *chooseR* algorithm with default parameters (https://github.com/rbpatt2019/chooseR)^18^.

For each resolution, we computed the adjusted Rand index (ARI) between unsupervised scRNA-seq clusters and author-curated cell-type clusters (i.e. ground truth) using the *adj*.*rand*.*index* (*fossil* R package, v 0.4.0)^64^. ARI is a measure of similarity between two data clusterings, adjusted for chance groupings. Across all the candidate resolutions evaluated, the maximal ARI between our unsupervised clusters and ground truth clusters was ∼0.8 and the resolutions at which the max ARI was observed was denoted *ARI*_*max*_ (**Fig 3B**). The imperfect cluster similarity here reflects differences in computational preprocessing across datasets and possible manual cluster refinement performed by authors of the original datasets. Nonetheless, this represents the maximal ARI that is achievable using the current unsupervised cluster approach and serves as a positive control to which all other cluster configurations were compared.

### Cell-type marker catalog

To generate a cell-type marker reference catalog, cell-type-specific markers were derived from eight diverse public scRNA-seq atlases (Tabula Muris^59^, Tabula Sapiens^60^, Cao 2019^3^, Cao 2020^57^, Pijuan Sala^24^, Tyser^16^, La Manno^27^ and Zeisel^58^) using the Wilcoxon DE method to identify DEGs across author-curated cell types (**Table 1**). All markers satisfying logFC > 0.5, AUROC > 0.95 and FDR < 1% were included. If less than 15 markers were identified per a cell-type using these criteria, the top N markers (ranked by logFC) with FDR < 1% were taken to ensure the minimum 15 markers per cell-type requirement was satisfied. These markers were then consolidated with cell-type-specific markers from PanglaoDB^21^ and CellMarkers^20^ to yield a cell-type marker reference catalog. No additional filtering was performed, resulting in many cell-types being represented by multiple gene sets from several independent sources. We justified this redundancy as a strength of the catalog, as co-enrichment of independent and coherent cell-type terms leads to higher confidence cell-type annotations. To visualize the catalog using a bipartite network, a gene × cell-type incidence matrix was generated using *graph*.*incidence* (*igraph* R package, v 1.2.6) and the network was visualized using *layout*.*auto* (*igraph*). Both human and murine cell-types are represented in this catalog. All cell-type markers used in this study have been made available in our scMiko R package.

### Cell-type annotation

The Miko score is a scaled cluster-level module score that adjusts for cell-to-cell gene expression variation and gene set size. To compute the Miko score, standardized module scores *Z*_*j*_ for each cell *j* must first be calculated by subtracting the mean expression of control features *Y*_*j*_ from the mean expression of gene set features *X*_*j*_, and then scaling the difference by the pooled standard deviation of the gene set and control features:

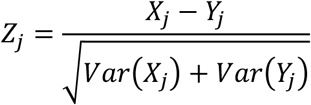

Following the approach taken by Tirosh and colleagues^65^ and implemented in *AddModuleScore* (Seurat), all analyzed features are binned based on averaged expression and control features are randomly selected from each bin. As a variance-corrected statistic, the standardized module score can be used as-is to compute single-cell level significance [*p* = Pr(> |Z|)]. However, in the absence of a gene set-size correction, module score comparisons between gene sets are invalid.

To correct for gene set size-dependencies, cell-level null standardized module scores *Z*_*null,j*_ are computed for randomly sampled gene sets that span over a range of different sizes (2-100 genes per gene set by default). Random gene set-specific *Z*_*null,j*_ scores are then aggregated for each cluster *k* to yield a cluster-level null standardized module score *Z*_*null,k*_:

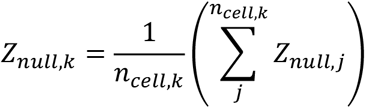

where *Z*_*null,k*_ and *Z*_*null,j*_ represent the null standardized module scores for a randomized gene set of a given size for cluster *k* or cell *j*, respectively, and *n*_*cell,k*_ represents the number of cells belonging to cluster *k*. The relationship between gene set size and null standardized scores is then fit using a polynomial spline:

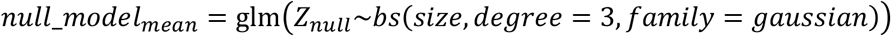

This null mean model is used to predict gene set size-adjusted null standardized scores 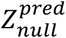. In theory, the expected value of 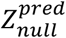 is 0 and we approximate it as such in our computational implementation. Separately, we calculate the observed variance in *Z*_*null,k*_, denoted *Var*(*Z*_*null,k*_), over a range of gene set sizes, and fit the relationship between gene set size and *Var*(*Z*_*null,k*_) using a gamma-family generalized linear model:

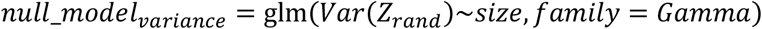

This null variance model is used to predict gene set size-adjusted variance of standardized scores 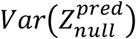.

Finally, to derive the gene set-size corrected Miko score, we aggregate standardized module scores *Z*_*j*_ for each gene set into cluster-level means:

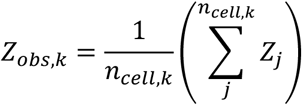

and center and scale *Z*_*obs,k*_ using gene set-size matched null mean 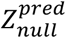 and variance 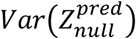 to yield the Miko score *M*_*k*_ for cluster *k*:

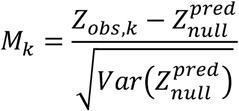

The Miko score is a cluster-level module score that is adjusted for gene set size-related spurious effects and cell-to-cell variability. This ensures the valid comparison of scores across differently sized gene sets, making it a valuable tool in marker-based cell annotation. Another property of the Miko score is that it can be handled as a Z statistic, thus facilitating p-value calculation and hypothesis testing:

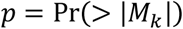

This facilitates cell cluster annotation based on which cell-type-specific gene sets are significantly active.

In addition to the Miko score, we propose two post-scoring filters which serve to fine tune which gene sets are considered enriched. The first is a coherence filter in which a positive correlation between component gene expression and the Miko score is enforced for a minimum fraction of component genes. The second is a frequent flier filter, which flags gene sets that exceed a minimum significance rate and represent gene sets that enrich across most cell clusters.

### Gene program discovery

<u>S</u>cale-free topology *<u>s</u>*hared nearest neighbor *<u>n</u>*etwork (SSN) analysis is a gene program discovery algorithm that groups genes based on co-expression similarity profiles and visualizes the network layout using a UMAP-based embedding. Features used for gene program discovery can be pre-specified using a variety of criteria, including minimum expression thresholds, high variability or deviance^66^, however in the current study we select features using a minimal expression criteria (expressing fraction > 0.5 within at least one cluster). The cell × gene expression matrix (transposed from the Seurat object) is then subject to principal component analysis [*RunPCA*(…, ndim = 50)] and the top components explaining >90% of the variance are used to construct a K-nearest neighbor graph *K* [*FindNeighbors*(…, k.param = 20)], from which a shared-nearest neighbor (SSN) graph *G* is constructed by calculating the neighborhood overlap (Jaccard Index) between every gene and its K-nearest neighbors. Adopting the framework from weighted gene correlation network analysis (WGCNA)^25,49^, a scale-free topology transform is then applied to the SNN graph by raising the SNN graph (gene × gene matrix) to an optimized soft-threshold power:

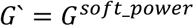

where *G*’ represents a scale-free topology-conforming SNN graph and is the adjacency matrix that will be used for downstream network construction. The optimal soft-threshold power used to derive *G*’ is identified by calculating the signed *R*^2^ statistic for the following relationship:

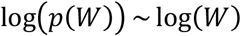

where *W* represents connectivity *w* discretized into *n* bins (default 20), and *p*(*W*) represents the proportion of nodes (i.e., genes) within the *W* bin. Connectivity *w*_*g*_ for gene *g* is calculated as row-wise sum of *G*:

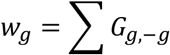

where *g* and −*g* represent the row and column indices corresponding to gene *g* and all genes except gene *g*, respectively. The soft threshold power is evaluated over a range of candidate values (default 1 to 5), and the optimal power is taken as the smallest power for which signed *R*^2^ < −0.9:

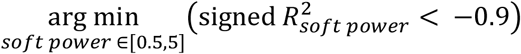

To visualize the transcriptomic network, the scale-free SNN graph *G*’ is embedded in a UMAP using *RunUMAP*(…, graph = *G*‘, umap.method = “umap-learn”). Network nodes represent individual genes, whereas network linkages represent *G*’ edges thresholded at a specified quantile (0.9 by default).

To identify gene programs from the scale-free SNN graph *G*’, Louvain clustering is performed. We identify the optimal clustering resolution using a ***nearest neighbor purity criterion*** which seeks to optimize the cluster consistency, or purity, within individual gene neighborhoods by maximizing the similarity of genes within programs compared to other programs (analogous to silhouette score^17^). For a candidate cluster resolution *ρ*, the gene-level purity score is defined as the proportion of genes within gene *g*’s neighborhood that belong to the most represented cluster within that neighborhood (**Fig S1**):

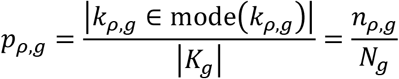

where *p*_*ρ,g*_ is the gene *g*’s purity at *ρ* resolution, the denominator *N*_*g*_ represents the cardinality (||), or size, of gene *g*’s K-nearest neighborhood *K*_*g*_ (20 by default), the numerator *n*_*ρ,g*_ represents the number of genes in gene *g*’s neighborhood that belong to the most represented cluster [i.e., majority cluster, mode(*k*_*ρ,g*_)] and *k*_*ρ,g*_ is a vector of cluster memberships for all genes belonging to gene *g*’s neighborhood. For each candidate resolution, gene-level purity scores *p*_*ρ,g*_ are then aggregated as means to yield the global purity score *P*_*ρ*_:

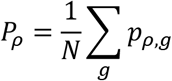

where *N* is the number of genes in the SSN graph. Finally, the optimal cluster resolution is the maximal resolution at which the target purity *P*_*target*_ (0.8 by default) is satisfied:

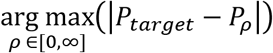

Possible purity scores range between 0 to 1. Neighborhoods in which genes belong to many different clusters are considered “impure” (low purity score) whereas neighborhoods in which genes belong to a single cluster are “pure” (high purity score). In general, higher cluster resolutions are associated with lower the purity scores, however we recommend using a target purity between 0.7 (more gene programs) and 0.9 (fewer programs).

To minimize spurious gene program associations, genes with low connectivity (i.e., low network degree) are pruned so that only hub-like genes are retained for downstream annotation and analysis. Here connectivity for each gene *g* is calculated as described above for *w*_*g*_, however in this case we use the scale-free SSN graph *G′* instead of *G*. Connectivity scores *w*_*i*_ are L2 normalized and those below a prespecified threshold (0.1 by default) are pruned.

### SSN performance evaluation

To benchmark the performance of SSN, gene program discovery was performed using SSN, independent component analysis (ICA) and non-negative matrix factorization (NMF) on eight public scRNA-seq data sets (**Table 1**). For each dataset, a common subset of genes that was expressed by >50% of cells in at least one cell cluster were used (typically ranging between 1000-4000 genes). ICA was performed using *RunICA*(…) implemented in *Seurat* (default parameters), and NMF was performed using *nnmf*(…, k = c(5, 10, 15), loss = “mse”, rel.tol = 1e-4, max.iter = 50) (*NNLM* R package, v 0.4.4). For NMF analysis, scaled gene expression values were truncated at zero. ***Graph modularity*** was compared between SSN graphs before (SN) and after (SSN) scale-free topology transformation using *modularity*(…) (*igraph* R package, v 1.2.6). ***GO*** g***ene set recovery*** was evaluated following the approach taken by Saelens and colleagues^31^, where the Jaccard similarity between observed (SSN, ICA, NMF) and known (GO) gene programs was calculated to yield an observed × known gene program similarity matrix. Then, for each known gene program (matrix column), the max column-wise Jaccard similarity score was taken, representing the best recovery achieved by the unsupervised gene program detection algorithm for that known gene program, and the best Jaccard indices averaged over all known programs yielded the overall recovery score. The overall recovery score was compared across gene program detection methods. To evaluate the extent of ***STRING protein-protein interaction enrichment*** in gene programs identified by each method, within-program interaction enrichment was performed using *get_ppi_enrichment*(…) (*STRINGdb* R package, v 2.0.2) and enrichment ratios were compared across gene program discovery methods^67^. Finally, we used the murine gastrulation scRNA-seq data set to benchmark the ***computing times*** required to run each method. The data set was subsampled to 1000, 10000, 25000, 50000 and 100000 cells and for each data subset, 500, 1000, 2500, 5000, and 10000 genes were used for gene program discovery. The run times, relative to SSN, as well as the absolute run times for SSN across different cell/gene count settings were reported.

### Gene set enrichment analysis

To functionally-annotated gene programs identified by SSN, ICA and NMF, we perform hypergeometric overrepresentation analysis using *fora* (*fgsea* R package, v 1.14.0)^68^. Annotated gene sets used for enrichment analyses included GO ontology (biological processes, cellular components, molecular function) and gene-set collections curated by the Bader Lab^69^.

### Data visualization

Unless otherwise specified, the *ggplot2* R package (v 3.3.5) was used for data visualization. scRNA-seq gene expression was visualized using *FeaturePlot* function (*Seurat*) or *DotPlot* function (*Seurat*). Venn diagrams were generated using either *ssvFeatureEuler* (*seqsetvis* R package, v 1.8.0) or *ggVennDiagram* (*ggVennDiagram* R package, v 1.1.4).

## Supplemental Figure

**Supplemental Figure 1.**
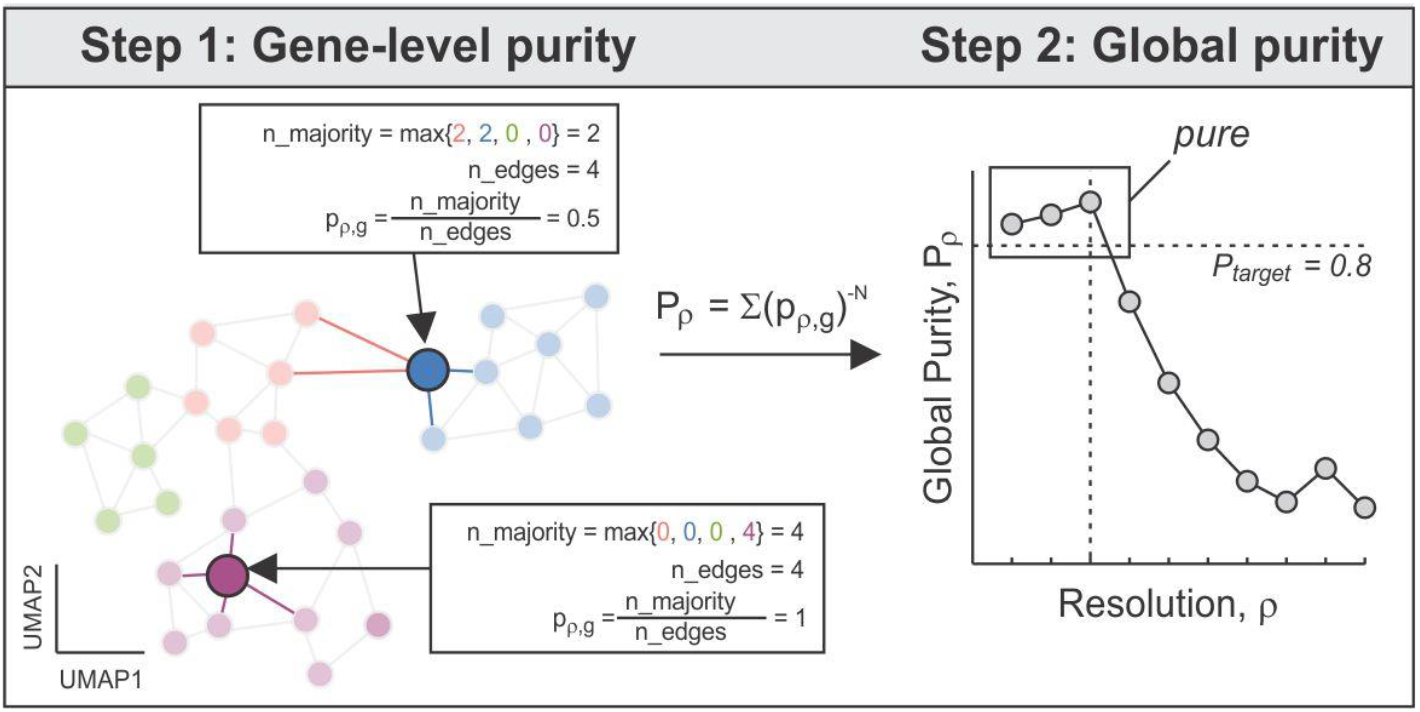
Schematic of nearest neighbor purity criterion. The nearest neighbor purity criterion seeks to optimize the cluster consistency, or purity, within individual gene neighborhoods by maximizing the similarity of genes within gene programs compared to other programs. In step 1 (*left*), for a given cluster resolution *ρ*, the gene-level purity score *p*_*ρ,g*_ is defined as the proportion of genes within gene *g*’s neighborhood that belong to the most represented cluster within that neighborhood. The gene-level purity scores *p*_*ρ,g*_ are then aggregated as means to yield the global purity *P*_*ρ*_. In step 2 (*right*), the optimal cluster resolution (*vertical dashed line, right*) is the maximal resolution at which the target purity *P*_*target*_ is satisfied (0.8 by default; *horizontal dashed line, right*).

**Supplemental Figure 2.**
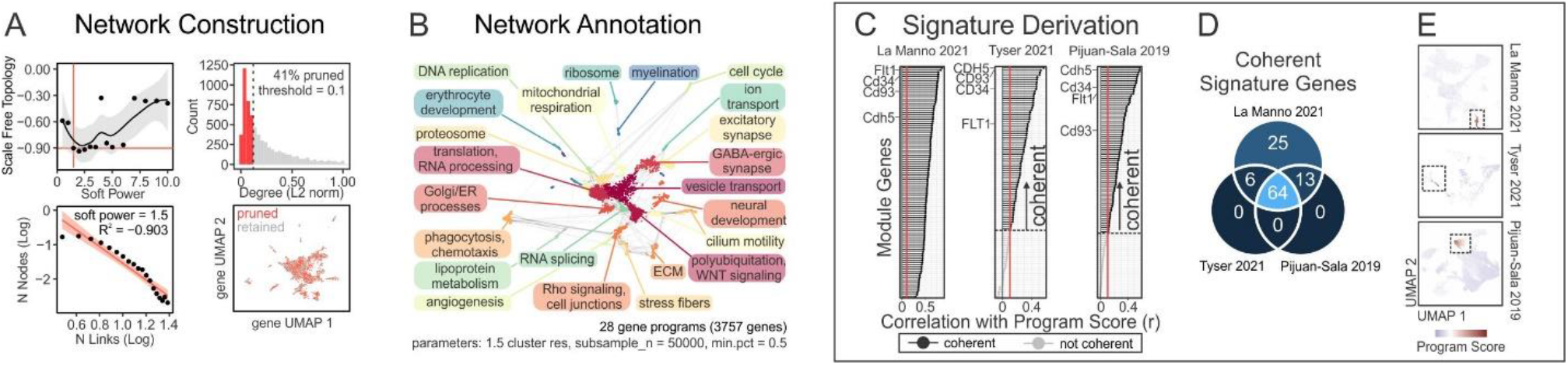
Application of SSN to identify robust angiogenesis-associated gene program. **(A-E)** Representative transcriptional network construction, annotation and applications using La Manno 2021 scRNA-seq data^27^. **(A)** Optimal soft power required for scale-free topology (*left column*; threshold = 0.9) and pruning of genes with low network connectivity (*right column*; threshold = 0.1). **(B)** Functional annotation of gene programs. **(C-E)** External validation and refinement of angiogenesis signature. **(C)** Correlation between angiogenesis program activity and expression of individual gene program genes across three independent scRNA-seq datasets. Genes that exceeded the coherence threshold (Spearman correlation > 0.1) were deemed coherent. (**D**) Venn diagram illustrating intersection between coherent gene sets determined in each scRNA-seq dataset. 64/108 genes (59%) were coherent in all scRNA-seq datasets. **(E)** Gene program activity of coherent angiogenesis signature specifically highlights the (hematogenic) endothelial population in all three scRNA-seq datasets.

## Acknowledgements

The authors thank members of our single-cell team, collaborators, and the Donnelly Sequencing Centre for technical assistance and/or helpful discussions. Our research was funded by grants from the Canadian Institutes of Health Research (J.M. and H.H.), Canada First Research Excellence Fund Medicine by Design Program (J.M.), and Donnelly Centre Home Research Fellow Fund (H.H.). N.M. was supported by the 2020 William Donald Nash Brain Tumour Research Fellowship.

## Author Contributions

Study design, data analysis, and interpretation: N.M., K.B., H.H.; Development of computational tools: N.M.; Manuscript writing: N.M. and H.H. with input from other authors; Project Conceptualization: H.H., N.M., and J.M.; Supervision: H.H., J.M.; Funding Acquisition: J.M. and H.H.

